# Data-driven characterization and correction of the orientation dependence of magnetization transfer measures using diffusion MRI

**DOI:** 10.1101/2023.10.05.561088

**Authors:** Philippe Karan, Manon Edde, Guillaume Gilbert, Muhamed Barakovic, Stefano Magon, Maxime Descoteaux

## Abstract

**Purpose:** To characterize the orientation dependence of magnetization transfer (MT) measures in white matter (WM) and propose a first correction method for such measures.

**Methods:** A characterization method was developed using the fiber orientation obtained from diffusion MRI (dMRI) with diffusion tensor imaging (DTI) and constrained spherical deconvolution (CSD). This allowed for characterization of the orientation dependence of measures in all of WM, regardless of the number of fiber orientation in a voxel. Furthermore, a first correction method was proposed from the results of characterization, aiming at removing said orientation dependence. Both methods were tested on a 20-subject dataset and effects on tractometry results were also evaluated.

**Results:** Previous results for single-fiber voxels were reproduced and a novel characterization was produced in voxels of crossing fibers, which seems to follow trends consistent with single-fiber results. Unwanted effects of the orientation dependence on MT measures were highlighted, for which the correction method was able to produce improved results.

**Conclusion:** Encouraging results of corrected MT measures showed the importance of such correction, opening the door for future research on the topic.

## 1| INTRODUCTION

The myelin sheath wrapping the axons that compose white matter (WM) is being linked to the progression of many neurodegenerative diseases and is also a key part of neurodevelopmental research. As a result, the search for myelin specific measures has become increasingly crucial. One promising avenue of magnetic resonance imaging (MRI) is magnetization transfer imaging (MTI) ^1,2^, which uses the distinct absorption lineshapes of free water and bound water in and around macromolecules to induce a magnetization transfer (MT) from the bound pool to the free pool. Indeed, an off-resonance radio-frequency (RF) pulse can be applied to saturate the magnetization of the bound pool, which then transfers this magnetization to the free pool, effectively reducing the measured signal. Typically, one can compute the magnetization transfer ratio (MTR) by comparing the images obtained with one or two single positive/negative frequency-offset pre-pulses (*S*_+_/*S*_*−*_) and without any MT preparation pre-pulse (*S*_0_):

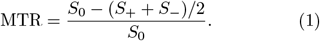

To some extent, this measure is sensitive to myelin ^3,4^ because of the macro-molecules that compose it, but is also influenced by other factors like inflammation ^5^, tissue components and tissue water content ^6^. Moreover, Helms et al. ^7^ proposed the MT saturation measure (MTsat) to be less impacted by T_1_ relaxation, flip angle and B1+ inhomogeneity effects:

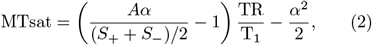

where *A* is the gain factor of the image for different repetition times (TR) and flip angles (*α*). Recently, Varma et al. ^8^ introduced an extension of the MT method called inhomogeneous magnetization transfer (ihMT), supposed to be more specific to myelin, unlike MT. Although the physical origin of the ihMT contrast was first unclear and up to debate ^9,10^, a recent effort from Alsop et al. ^11^ helped to clarify the questions. They explain that ihMT isolates the contribution of the dipolar order from the MT phenomena. This results in ihMT being only sensitive to bound water where protons undergo dipolar interactions at long dipolar order relaxation time. Lipid membranes correspond to such environment and are a major component of myelin, hence the specificity of ihMT to it. The ihMT ratio (ihMTR) is calculated from the images obtained with single positive/negative frequency-offset pre-pulses (*S*_+_/*S*_*−*_), with dual positive-negative/negative-positive alternating-offset pre-pulses (*S*_+*−*_/*S*_*−*+_) and without any MT preparation pre-pulse (*S*_0_):

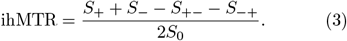

Similarly to the MTsat measure, the ihMT saturation (ihMTsat) is calculated as ^12,11^:

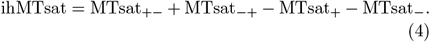

As eluded previously, dipolar order affects both MT and ihMT measures. It is well known that dipolar interactions depend on the angle (*θ*_n_) between the vector connecting the dipoles and the direction of the main magnetic field **B**_0_, following a (3 cos^2^ *θ*_n_ *−* 1)*/*2 relationship.

It has been observed that parameters of the quantitative MT binary spin bath (BSB) model ^13^ are correlated to the orientation of white matter fiber bundles ^14,15^. Pampel et al. ^16^ further explored this phenomenon and proposed a novel absorption lineshape for the BSB model that takes into account the orientation of fiber bundles. More precisely, they modelled the myelin sheath as lipid bilayers wrapped around an axon in a cylindrical way. This implies that the vector connecting the dipoles is always normal to the cylinder, as shown in figure 1, hence the notation *θ*_n_.

**FIGURE 1.**
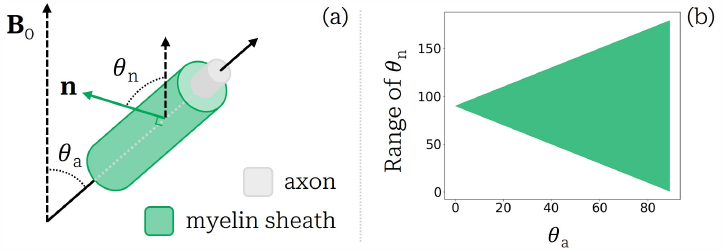
(a) Illustration of the angles in play in the cylindrical model for the myelin sheath. The angle between the main magnetic field **B**_0_ and the direction of the axon is *θ*_a_, while the angle between **B**_0_ and a vector normal to the myelin sheath (**n**) is *θ*_n_. Adapted from Girard et al. ^17^. (b) Range of values that *θ*_n_ can take depending on *θ*_a_.

More recently, similar research has been done by Girard et al. ^17^ on ihMT, once again reporting an angular dependency of the myelin lineshape. Furthermore, Morris et al. ^18^ measured the orientation dependence of a few parameters, including the ihMTR, in a phospholipid bilayer, and found a relationship that corresponds well with dipolar order. Using their results, they also simulated the variation of ihMTR with respect to the angle (*θ*_a_) between a fiber bundle and the main magnetic field, in the case of a cylindrical myelin sheath. However, an analysis on *in vivo* data from a multi-subject dataset showed a different behavior of ihMTR in this regard ^19^. Nevertheless, this study also exhibits the dependency of MTR, which seems to follow a trend similar to the simulated ihMTR from Morris et al. ^18^ and the local dipolar field calculated by Girard et al. ^17^.

Diffusion MRI, although not known for being very specific to myelin, is the modality of choice when it comes to reconstructing the WM fiber population orientations in each voxel. The simplest way to do so is using diffusion tensor imaging (DTI) ^20^ to get the principal eigenvector of the diffusion tensor. Although this is not robust in voxels of crossing fibers ^21^, which represent a large fraction of the voxels in the brain ^22,23^, it still prevails in single-fiber voxels. For the case of crossing fibers, one may consider the use of a compartment model ^24,25^, or the widely used constrained spherical deconvolution (CSD) model ^26,27^. The latter provides a fiber orientation distribution function (fODF) representing the orientational disposition of fibers in each voxel. It is worth noting that Pampel et al. ^16^ used this approach, while ^19^ chose the principal eigenvector from DTI.

In this work, we further explore the orientation dependence of MT measures using state-of-the-art diffusion MRI techniques. We first try to replicate results from past studies in single-fiber voxels for MTR and ihMTR, as well as MTsat and ihMTsat, by computing the mean of these measures with respect to the angle *θ*_a_. We then study the behavior of such measures in voxels of multiple fibers crossing, using fODF information obtained from CSD. This characterization also leads to a first attempt at correcting these MT measures according to the WM fiber orientation in each voxel. Indeed, none of the previous studies proposed an angular correction for MTR or ihMTR measures. Impacts of the corrections on contrasts as well as tractometry are discussed and promising results set the table for future development.

## 2.| METHODS

### 2.1| Dataset acquisition

The dataset used in this study comes from Edde et al. ^28^. In summary, twenty healthy adults were scanned on a clinical Ingenia 3T MRI scanner (Philips Healthcare, Best, Netherlands) with a 32-channel head coil. The 33 minutes acquisition was repeated 5 times over 6 months for each participant, for a total of one hundred sessions. These acquisitions include 3D T1-weighted images, multi-shell diffusion-weighted images (DWI) and ihMT images. More details can be found in Edde et al. ^28^.

### 2.2| Processing

DWI and T1w images were processed with Tractoflow ^29^ to denoise, correct and register the data (see Theaud et al. ^29^ for further details). Tractoflow also generated the fODFs using CSD from the Python library DIPY^30^, and extracted the number of fiber orientation (NuFO), the fODF peaks and peak amplitudes (also called peak values) using the *scil compute fodf metrics*.*py* script from the Python library Scilpy (https://github.com/scilus/scilpy). From these, peaks fractions were calculated by normalizing the peak values per voxel. DTI eigenvectors and eigenvalues, as well as DTI measures such as the fractional anisotropy (FA), were also computed by the pipeline using only b-values equal to or below 1200 s/mm^2^. Moreover, Tractoflow generated a tissue segmentation from the T1w images and a whole-brain tractogram from the fODF map and the tracking masks. See Edde et al. ^28^ and Theaud et al. ^29^ for more information on this pipeline. Some bundles of interest were also extracted from the tractograms using a multiatlas multi-parameter version of RecoBundles ^31^ called RecoBundlesX ^32^.

As explained in Edde et al. ^28^, the ihMT images were processed using a combination of multiple tools (FSL, ANTs ^33^, SCIL ihmt flow pipeline script https://github.com/scilus/ihmt_flow). We refer to Edde et al. ^28^ for a complete description of the processing of MT and ihMT measures, which were generated as described in Helms et al. ^7^ and Varma et al. ^8^ respectively. One notable difference with Edde et al. ^28^ is that the ihMTsat measure ^12^ is now computed instead of the ihMTdR1sat.

### 2.3| Single-fiber characterization

The first characterization step is the definition of orientation dependence of MT measures in single-fiber voxels. As shown in step 1 of figure 2, such voxels are selected by taking the intersection of three masks: a WM tissue mask, a mask where the FA is greater than a threshold, and a mask where the NuFO equals 1. For each voxel in this selection mask, we compute the angle *θ*_a_ separating the main magnetic field **B**_0_ and the direction of the principal eigenvector **e**_1_ of the diffusion tensor, obtained through DTI. For the sake of keeping *θ*_a_ between 0 and 90 degrees, angles greater than 90 degrees are subtracted to 180, using the fact that the diffusion tensor is symmetrical. The voxels are then put in angle bins of a certain width (Δ^*°*^), and the number of voxels in each bin is saved. The mean of the chosen measure is computed for each bin and can be plotted afterwards as a function of *θ*_a_, as shown in step 2 of figure 2. Moreover, bins that have under 30 voxels are not plotted, as they do not contain enough data to be representative (this is also true for the following section).

**FIGURE 2.**
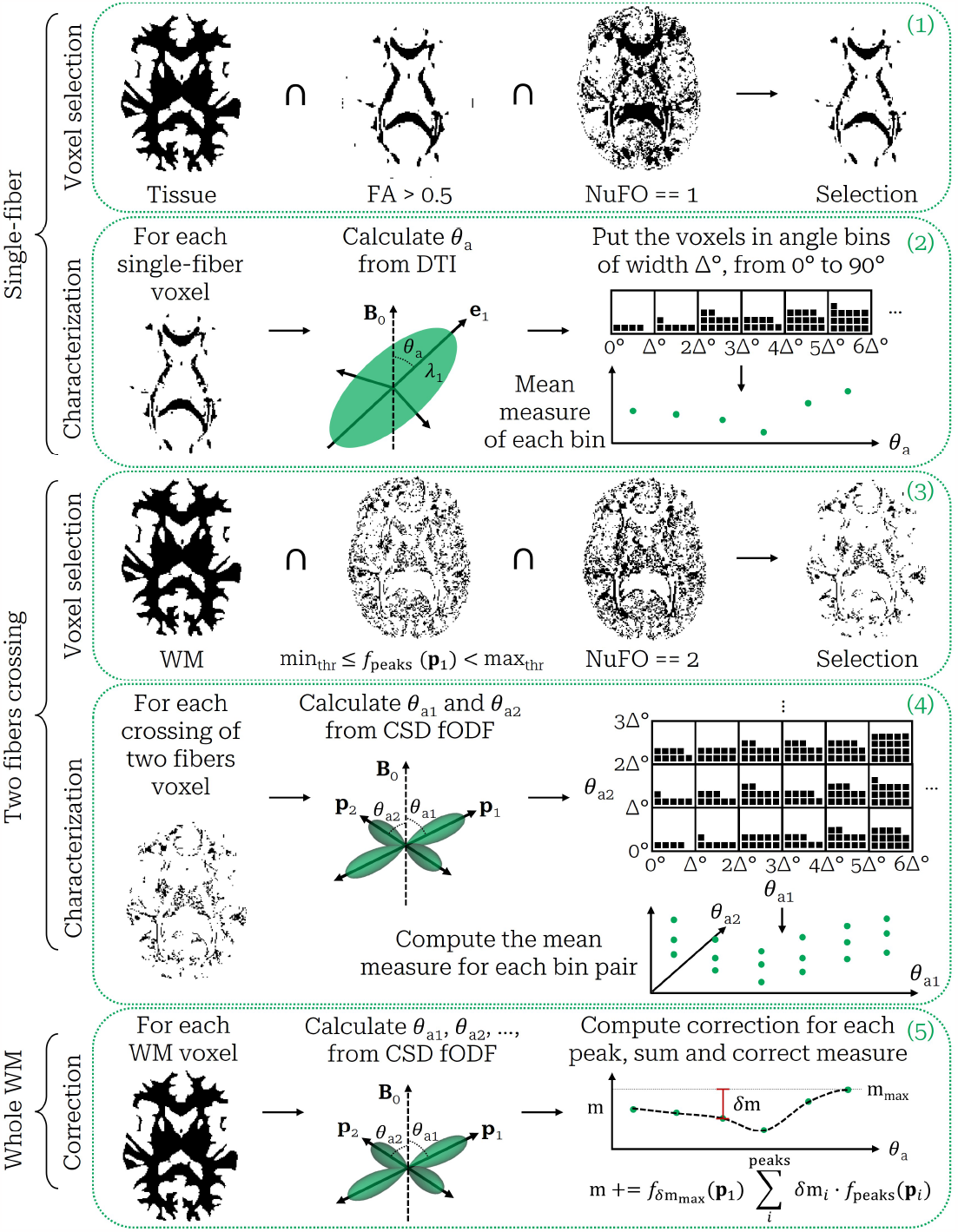
Summary of the characterization and correction methods. In the first step (1), the single-fiber selection mask is created as the intersection of a tissue mask, a FA threshold mask and a NuFO equals 1 mask. In the second step (2), the angle *θ*_a_ between **B**_0_ and the principal eigenvector **e**_1_ of the diffusion tensor is calculated for each selected voxel, and put in angle bins of width Δ^*°*^. The measures are then averaged for each bin and plotted as a function of *θ*_a_. In the third step (3), the two-fibers crossing selection mask is created as the intersection of a WM mask, a peaks fraction threshold mask and a NuFO equals 2 mask. In the fourth step (4), the angle *θ*_a_ between **B**_0_ and each of the two fODF peaks **p**_1_ and **p**_2_ is calculated for each selected voxel, and put in a 2D matrix of angle bins of width Δ^*°*^. The measures are then averaged for each matrix element and plotted as a function of *θ*_a1_ and *θ*_a2_. In the fifth step (5), the angle *θ*_a_ of each fODF peak (up to five per voxel) in the WM is used to compute the correction from the fitted characterization curve. The total correction for each voxel is computed as the sum of each peaks correction weighted by their fraction (*f*_peaks_) and the maximum amplitude factor 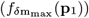of the voxel.

This characterization method assumes an homogeneous distribution of myelin around WM, which is not necessarily true, as pointed out by Morris et al. ^19^. Indeed, an extension of COMMIT allowing myelin streamline decomposition ^34^ revealed variations of myelination between WM bundles. Such assumption could result in overestimation of the measures mean at certain angles, especially for ihMT. For instance, the corticospinal tract (CST), which axons are larger than in other tracts ^35^, could be highly myelinated and thus increase the measures mean at low *θ*_a_ angles, as CST is roughly aligned with **B**_0_. With that in mind and taking inspiration from Kauppinen et al. ^36^, we also try a characterization method using a corpus callosum (CC) mask instead of a whole WM mask. The CC is a WM fiber bundle spanning across a large portion of the brain, therefore encompassing a wide variety of *θ*_a_ angles. This mask should allow the method to evaluate angles from 0 to 90 degrees, while assuming better homogeneity of the myelin distribution.

### 2.4| Crossing fibers characterization

The characterization of the orientation dependence of MT measures in voxels containing two fibers crossing is similar to the single-fiber characterization, with a few adjustments presented in steps 3 and 4 of figure 2. The selection mask is calculated as the intersection of a WM mask, a mask where the NuFO equals 2 and a mask selecting a certain range of peak fraction *f*_peaks_(**p**) for the first peak (**p**_1_). This last mask allows for the selection of a particular set of proportions for the two fibers in the voxel, as their fractions sum to one. Next, we compute the angle of the two fibers (*θ*_a1_ and *θ*_a2_) for each voxel in the selection mask, this time using the fODF peaks (**p**_1_ and **p**_2_) as shown in step 4 of figure 2. Angles greater than 90 degrees are again subtracted to 180, justified by the fact that the fODF is symmetrical. Every selected voxel is now described by two angles, that are both binned the same way as in section 2.3. This creates a 2D matrix of angle bins, with the axes being *θ*_a1_ and *θ*_a2_, covering all combinations of two fibers orientations. Since it is impossible to differentiate **p**_1_ from **p**_2_, each offdiagonal opposite component ((*i, j*), (*j, i*)) of the matrix are averaged together, yielding a symmetric matrix at the end. The number of voxels in each matrix element is also preserved by summing (*i, j*) and (*j, i*) in the off-diagonal elements. Finally, the mean of the chosen measure is computed for each matrix element, rendering a 3D plot of the orientation dependence of the measure. The diagonal of the matrix can also be plotted as a 2D plot resembling the previously introduced single-fiber plot, as both *θ*_a1_ and *θ*_a2_ are equal on the diagonal.

The 2D plot can be compared to the singlefiber plot to get a sense of the behavior in multiple fiber orientations voxels. In particular, the maximum amplitude *δ*m_max_ of the curve, defined as the separation between the minimum and the maximum, can be computed for various peak fractions, namely *f*_peaks_(**p**_1_) *∈ {*[0.5, 0.6[, [0.6, 0.7[, [0.7, 0.8[, [0.8, 0.9[*}*. A fit of these four points results in a continuous function, *δ*m_max_(*f*_peaks_(**p**_1_)), that can then be normalized by the maximum amplitude of the single-fiber curve, leading to a factor linking the amplitude of any crossing fibers curve to the reference single-fiber curve.

Furthermore, the characterization of voxels containing three fiber orientations can also be performed with the same method by simply adding another angle *θ*_a3_ for the third peak (**p**_3_). As this leads to a 4D visualization, only the diagonal of the 3D matrix is plotted in this case. Since it is less likely to find three fibers with the same angle *θ*_a_, a large bin width is needed (30 degrees).

### 2.5| Correction method

The correction method is mostly based on the results of the single-fiber characterization, using either the whole WM mask or the CC mask. However, it can also take into account the maximum amplitude factor computed from the crossing fibers analysis. As presented in step 5 of figure 2, the single-fiber orientation dependence plot is fitted using a polynomial fit of degree 10, allowing a continuous description of the dependence at every angle. The degree of the polynomial fit was chosen by testing multiple values and evaluating the better fit throughout subjects. Then, for each fODF peak, in every voxel of the WM mask, the angle *θ*_a_ is calculated. The idea is to bring all the measures to the maximum value of the fitted curve (m_max_), effectively removing the effect of orientation. Thus, for each calculated angle, the difference (*δ*m) between m_max_ and the fit at *θ*_a_ is computed. All the differences of each peaks in a voxel are then summed up, weighted by the peaks fractions (*f*_peaks_(**p**)) and an optional maximum amplitude factor 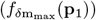, written as:

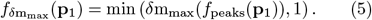

The corrected measure is given by this summed correction added to the original measure:

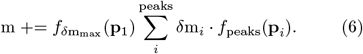

The Python code used for characterization and correction is available at https://github.com/karanphil/mtdiffusion/tree/devscript.

### 2.6| Impacts of correction on tractometry results

To further study the possible outcomes of correcting MT measures, we compare the corrected tractometry measures with the original ones presented by Edde et al. ^28^. Indeed, using the SCIL tractometry flow pipeline (https://github.com/scilus/tractometry flow ^37^), we compute both the bundle-average and track-profile ^38^ of each extracted bundles, for our four MT measures. A description of the bundles is available at https://high-frequency-mri-database-supplementary.readthedocs.io/en/latest/results/measure.html#af, as well as all original results from ^28^. Note that the middle cerebellar peduncle (MCP) was not present in this previous study, but is now added in the current work. Each bundle is separated into ten equidistant sections, resulting in ten-point track-profiles.

## 3| RESULTS

### 3.1| Single-fiber characterization

Figure 3 presents a first glance at the results of singlefiber characterization with MTR and ihMTR for subject 2, session 4. The voxel count, displayed as a grey-scale colormap of the markers, shows an increasing number of voxels with respect to the angle *θ*_a_. The figure also illustrates very different behaviors of orientation dependence for MTR and ihMTR. Indeed, MTR is maximal when the fibers are perpendicular to **B**_0_ (90 degrees) and minimal at around 40 degrees, while ihMTR is maximal when the fibers are parallel to **B**_0_ (around 0 to 10 degrees) and minimal when fibers are perpendicular to **B**_0_ (90 degrees). The following figures present various aspect of the single-fiber characterization. Note that each following plot uses the same pattern of circles for original measures and squares for corrected measures, as well as a dashed line for the polynomial fit. Furthermore, the variability of the characterization is assessed in Supporting Information and the reproduced results from Morris et al. ^19^ are presented. With these insights, all the following single-subject results come from subject 2, session 4, with a FA threshold of 0.5, a bin width of 1 degree for single-fiber voxels, a bin width of 10 degrees in the case of voxels of crossing fibers for increased voxel count, and only MTR and ihMTR.

**FIGURE 3.**
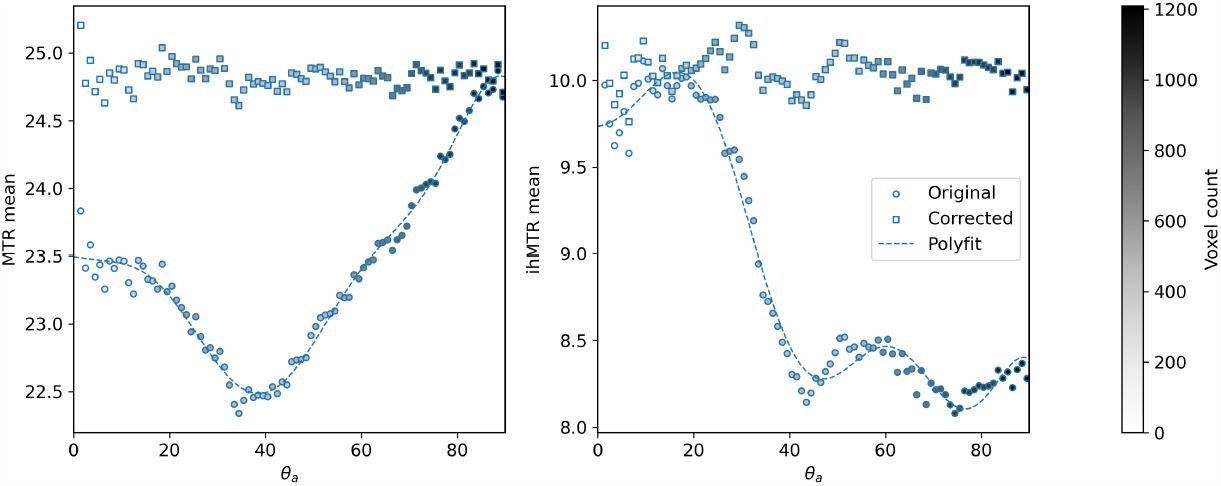
Mean MTR and ihMTR with respect to the angle *θ*_a_. The grey-scale colormap of the markers shows the voxel count per bin. The circle markers show the original measures, while the square markers represent the corrected measures. The dashed line is the polynomial fit on the original measures.

#### 3.1.1| Localization of angle bins in the single-fiber selection mask

The first row of figure 4 shows the distribution of angle *θ*_a_ for single-fiber voxels in the brain, using a bin width of 1 degree. The angle *θ*_a_ varies smoothly across the different structures of WM. It is worth noting that most of the low angle bins (0 to 40 degrees) are located in the CST, while a majority of the medium to high angle bins (40 to 90 degrees) are located in the CC, although these bins present more variable locations.

**FIGURE 4.**
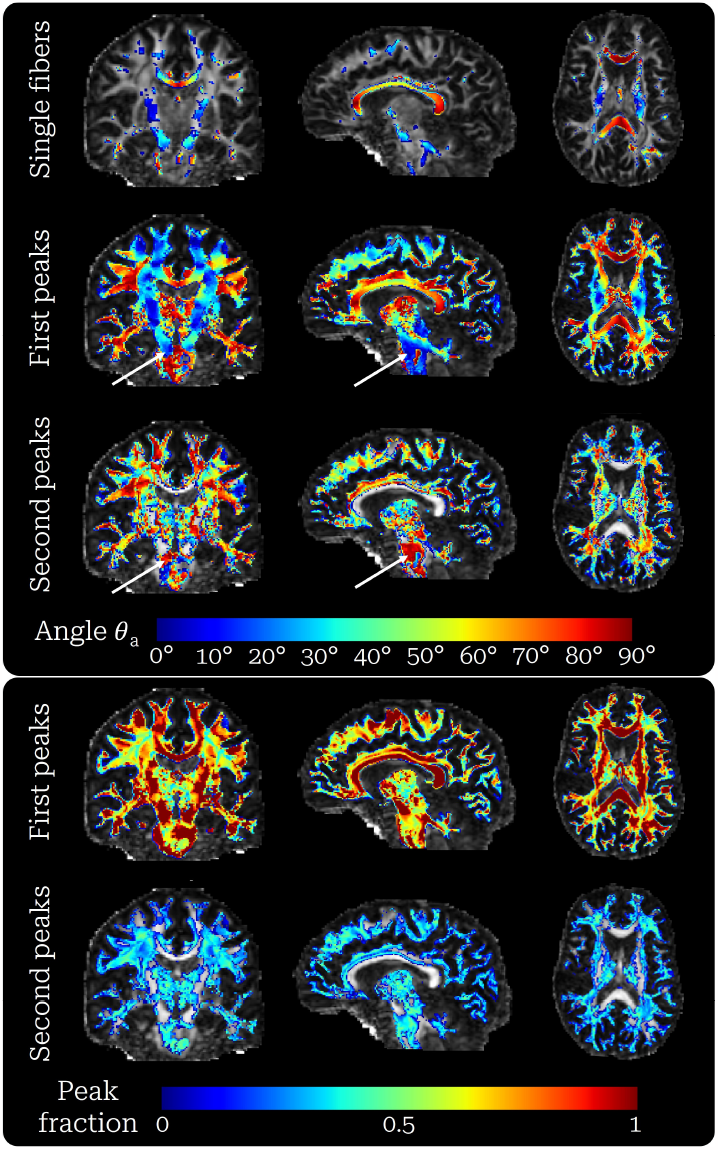
Distribution of angles *θ*_a_ and peak fractions across the brain. First row: angle *θ*_a_ of only the single-fiber voxels, extracted from the single-fiber method. Second and third rows: angle *θ*_a_ of the first and second peaks extracted from the crossing fibers method, respectively. Single-fiber voxels from the first row are included in the first peaks row. The white arrows highlight the crossing of the CST and the MCP. Fourth and fifth rows: fractions of the first and second peaks, respectively.

#### 3.1.2| Corpus callosum versus whole white matter

Figure 5 shows undeniable differences between the whole WM and the CC characterization curves. Indeed, the curves obtained in the CC have lower mean measures, from 0 to about 40 degrees, especially for ihMTR. Moreover, these curves still follow similar trends even if their amplitudes differ. The rest of the curves at angles higher than 40 degrees are very alike, especially for MTR which shows the exact same trend.

**FIGURE 5.**
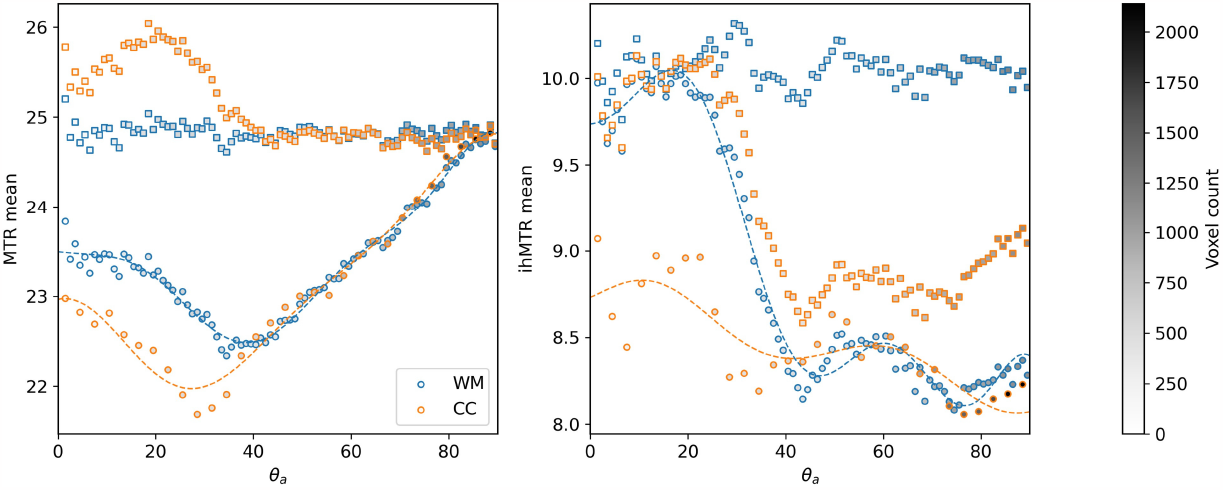
Mean MTR and ihMTR with respect to the angle *θ*_a_, for a characterization of the whole WM (WM) and a characterization limited to the corpus callosum (CC), both as circle markers. The dashed lines show there respective polynomial fit. Note that the bin width for the CC characterization is 3 degrees, to compensate for the lower amount of voxels. The square markers show the corrected measures of the whole WM when using either the WM or the CC characterization.

#### 3.1.3| Localization of angle bins in the whole white matter

Figure 4 shows the orientation with respect to **B**_0_ of the first and second peaks for the whole white matter. This allows for a better understanding of the localization of the angle *θ*_a_, color-coded as a heat-map, for the different anatomical WM structures or bundles of the brain. The figure highlights the continuous and smooth variation of orientation of the peaks, especially for the dominant peak. Indeed, holes in the single-fiber images are nicely filled when adding the first peak extracted from the fODFs. The bottom part of the figure also gives an idea of the proportion of the two peaks in each voxel, where the peak fraction is again color-coded as a heatmap from 0 (no peak) to 1 (only peak). Regions that appear in blue-green in both rows present crossings of two fibers with similar populations.

### 3.2| Crossing fibers characterization

Figure 6 presents a 3D visualization of the results from the characterization of voxels with two crossing fibers. The orientation dependence in such voxels appears to follow roughly the same patterns as from the single-fiber characterization. Indeed, when fixing either fiber to a given angle, the 2D curve of the other fiber resembles the curve of figure 3, weighted by the amplitude of the first fiber. Thus, the orientation dependence of crossing fibers seems to result from the linear combination of the orientation dependence of the underlying fibers. However, we observe that the trend is a bit different when one of the fibers is oriented at the maximum of the curves from figure 3 (80 to 90 degrees for MTR, 0 to 10 degrees for ihMTR). The maximum MTR becomes at 0 to 10 degrees and the maximum ihMTR becomes at 80 to 90 degrees, for the other fiber. In both cases, the highest measure occurs when one fiber is parallel to **B**_0_ and the other fiber is perpendicular to it.

**FIGURE 6.**
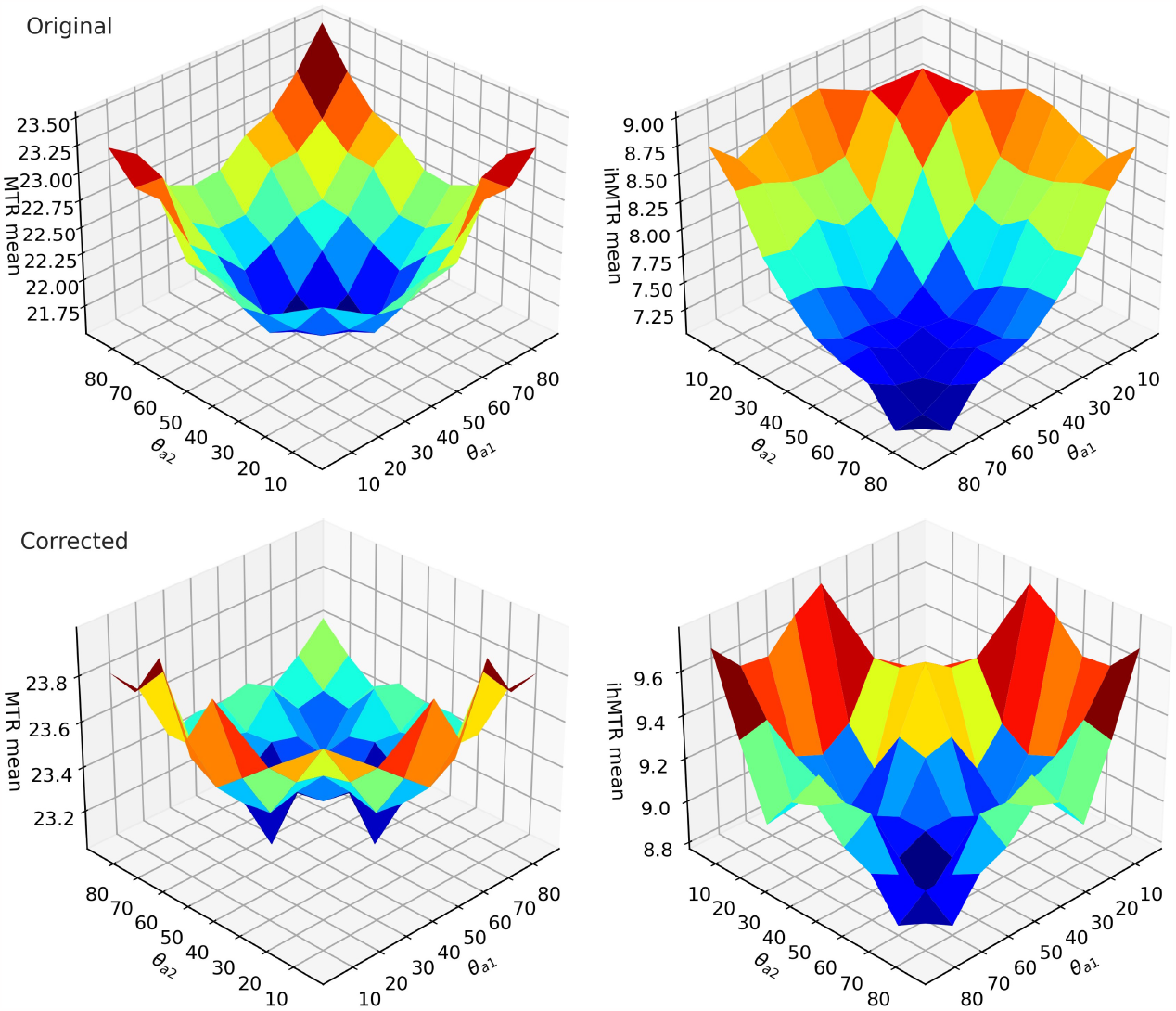
Mean MTR and ihMTR with respect to the angles *θ*_a1_ and *θ*_a2_, in the case of two crossing fibers with peak fractions between 0.5 and 0.6. Note that there is no mean measure when both fibers are at low angles (between 0 and 20 degrees), since this bin combination does not pass the minimal 30 voxel count. The first and second rows show the original and corrected measures, respectively. A jet map is used to help seeing the topology of the 3D surfaces.

Figure 7 illustrates the case where both fibers have the same orientation with respect to **B**_0_, for various ranges of peak fractions. Thus, the blue markers directly come from the diagonal of figure 6. Although the low angle points are missing, we still observe that the curves follow the same trend as the single-fiber characterization of figure 3. However, the maximum amplitude (*δ*m_max_) is not constant for each peak fraction. Indeed, the maximum amplitude decreases when the fraction of peak_1_ (*f*_peaks_(**p**_1_)) decreases, as presented in the subplot of figure 7. This relationship between the peaks fraction and the maximum amplitude of the curves is captured by the fitting function *δ*m_max_(*f*_peaks_(**p**_1_)) and can be used later on to account for peak fraction in the correction method. Since the maxima of the mean ihMTR are not available, this cannot be done efficiently for ihMT measures. Note that due to its low voxel count, the red point (**p**_1_ fraction between 0.8 and 0.9) was not taken into account by the fitting function, while all the other points align almost perfectly with the linear function. The figure in case of three fibers crossings is available as Supporting Information, providing equivalent information as figure 7. Even though the bins are very large (30 degrees), it is still possible to recognize the previously seen trends of MTR and ihMTR as well as the flattening effect of correction, especially on the most common peak fractions (0.3 to 0.5).

**FIGURE 7.**
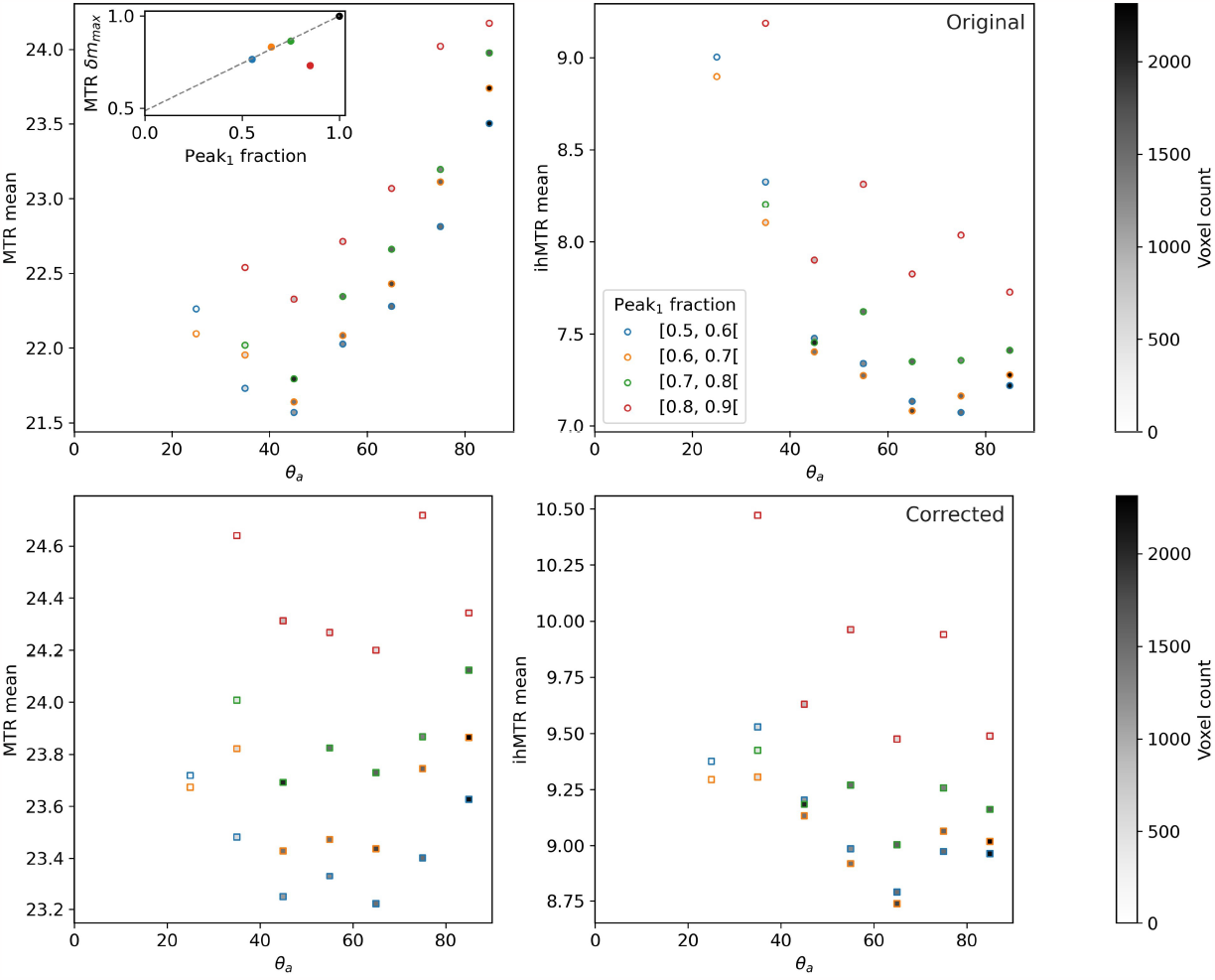
First row: original mean MTR and ihMTR with respect to the angle *θ*_a_, when angles *θ*_a1_ and *θ*_a2_ are equal, for various ranges peak fractions in the case of two crossing fibers. In other words, this is the diagonal of the matrices plotted in figure 6, but for different peak fractions. On the top left of the MTR plot is a subplot of the MTR *δ*m_max_ as a function of the fraction of the first peak. The grey dashed line shows the linear fit used in equation 5. Note that the MTR *δ*m_max_(*f*_peaks_(**p**_1_)) values were computed as the amplitude between the 40 to 50 degrees and 80 to 90 degrees bins, because the 40 to 50 degrees bin showed more reliability than the 30 to 40 degrees bin across subjects and sessions, having a higher voxel count. Second row: corrected mean MTR and ihMTR for the same situation as the first row.

### 3.3| Correction results

Figure 8 presents examples for which the orientation dependence of MTR and ihMTR is visible to the eye. The first row of these figures shows the locations of the absolute minimum and maximum of the characterization curves for MTR and ihMTR. The second row highlights regions where a difference in contrast is visible and most probably caused by the orientation dependence of the measures. Indeed, these regions agree with the locations of the first row and are often part of the same bundle or part of two major bundles such as the CC and CST. The third and fifth rows of figure 8 show the results of the correction method on MTR and ihMTR, using the whole WM or the corpus callosum characterization, respectively. While the two might look similar to the eye, the difference maps of the fourth and sixth rows suggest otherwise. For MTR, the correction using the CC characterization is more aggressive, as it occurs in high intensity across a good part of the WM. Conversely, the whole WM method is the more intense one for ihMTR, correcting strongly almost any WM voxels that do not have WM fibers parallel to **B**_0_. In either case, the correction method is able to remove the visual differences due to the orientation dependence, while keeping other details, especially for MTR.

**FIGURE 8.**
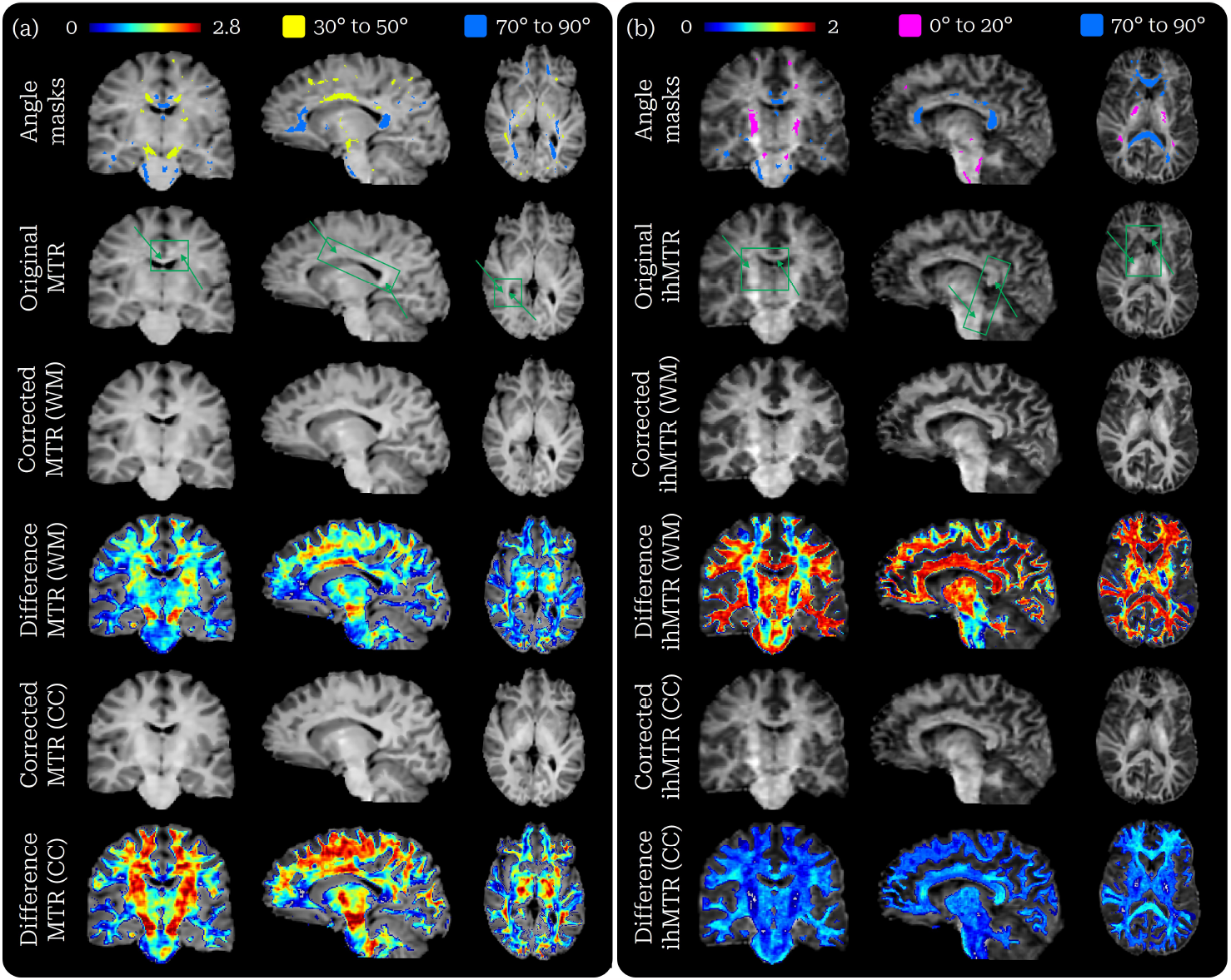
First row: masks of the 0 to 20 degrees (pink), 30 to 50 degrees (yellow) and 70 to 90 degrees (blue) bins. Second row: original MTR and ihMTR with green boxes and arrows highlighting regions where the orientation dependence is visible. Third row: corrected MTR and ihMTR using the whole WM (WM) characterization. Fourth row: difference map between the previous corrected and original MTR and ihMTR (in jet colormap). Fifth row: corrected MTR and ihMTR using the corpus callosum (CC) characterization. Sixth row: difference map between the previous corrected and original MTR and ihMTR.

Figure 3 shows the characterization curves of corrected MTR and ihMTR compared to the original ones, highlighting the removal of the orientation dependence for single-fiber voxels across white matter. Figures 6 and 7 also present a comparison of the corrected and original measures in the case of two WM fibers crossing, as a 3D plot and the diagonal version of it. The later shows a reduction of the orientation dependence regardless of the peaks fraction, as the curves become flatter after correction, especially for MTR. This is also visible on the 3D plots, which also highlight a limitation of the correction when one fiber is parallel to **B**_0_ and the other fiber is perpendicular to it. In such case, the measures means become higher than the rest of the possible fiber configurations. However, it is important to note that the range of variation in the corrected plots (*∼* 2) is much smaller than the range of the original plots (*∼* 1). The figure in case of three fibers crossings, available as Supporting Information, also demonstrates the ability of the method to properly correct such voxels.

The results of the whole WM correction using the corpus callosum characterization are compared to the whole WM method in figure 5. We show that the CC method is not able to properly remove the whole WM orientation dependence on both MTR and ihMTR. In the case of MTR, low angles are over-corrected, while the high angles are under-corrected for ihMTR.

### 3.4| Impacts of correction on tractometry results

Figure 9 compares original results from Edde et al. ^28^ of the mean MTR and ihMTR per bundle with the results obtained after correction. For both MTR and ihMTR, the observed trends between bundles follow roughly the same patterns before and after correction, with the exception of the CST. Indeed, the mean MTR of the CST is slightly increased with respect to the other bundles, when comparing the original and corrected measures. In the case of ihMTR, CST had by far the highest mean before correction and sustained a noticeable decrease with respect to other bundles after correction, while still remaining one of the highest ihMTR bundle. Overall, all bundles saw their mean MTR and ihMTR increase, except for the ihMTR of the CST. Moreover, the variability of the mean measures between sessions and subjects appears similar before and after correction. The same conclusions can be drawn from MTsat and ihMTsat measures, as shown in Supporting Information.

**FIGURE 9.**
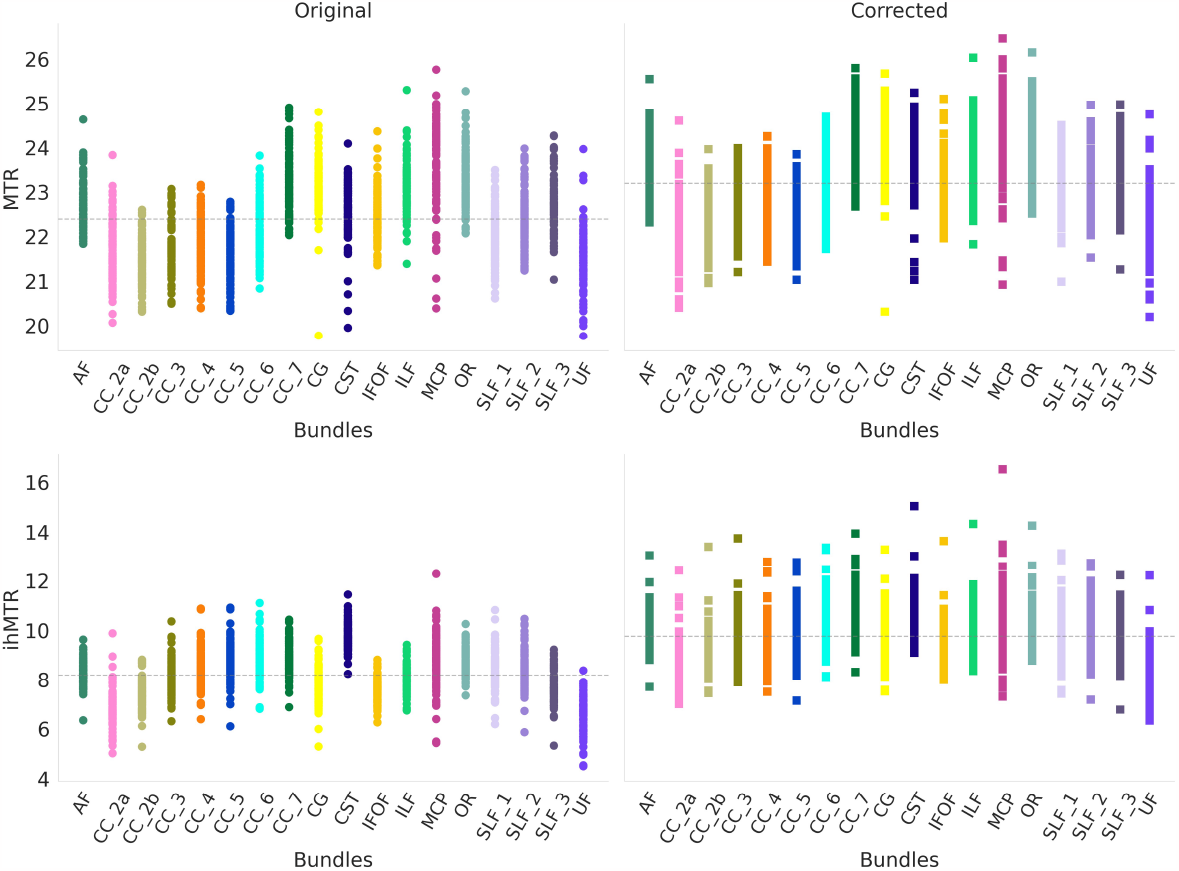
Mean MTR (top row) and ihMTR (bottom row) of each selected bundles, for each subjects and each sessions. Again, the circle markers correspond to original measures (on the left), while the corrected measures are represented by square markers (on the right). The dashed grey lines represent the mean value of all bundles, allowing for an easier comparison of changes between bundles. The left and right parts of bundles are averaged together.

Figure 10 showcases examples of the impacts of correction on some track-profiles, while also roughly illustrating the orientation of said tracks with respect to **B**_0_. On one hand, the CST is mostly aligned with **B**_0_ and presents only small deviations in trend for both MTR and ihMTR, but has increased measures after correction. On the other hand, the CC 3 possesses a wide range of orientations, from the central sections (5, 6) that are perpendicular to **B**_0_, to the endpoint sections (1, 10) which are parallel to it. As for its track-profile, various effects of the correction are observed. First, the MTR in central sections 5 and 6 is practically untouched, while endpoint section 1, 2, 9 and 10 are moderately increased. The biggest change comes from sections 3, 4, 7 and 8, in which the MTR profile is increased drastically. Second, the ihMTR shows similar changes as MTR, except for the central sections that are now greatly increased as well. In addition to the CST and CC 3 profiles, figure 10 presents the profile of the arcuate fasciculus (AF), which is mostly perpendicular to **B**_0_ from sections 5 to 10 and bends in such a way that section 3 is roughly aligned with the main magnetic field. Sections 5 to 10 of the AF, as well as section 1, show constant low increase for MTR and high increase for ihMTR. Moreover, the gap between the original and corrected profiles increases for MTR and decreases for ihMTR around section 3.

**FIGURE 10.**
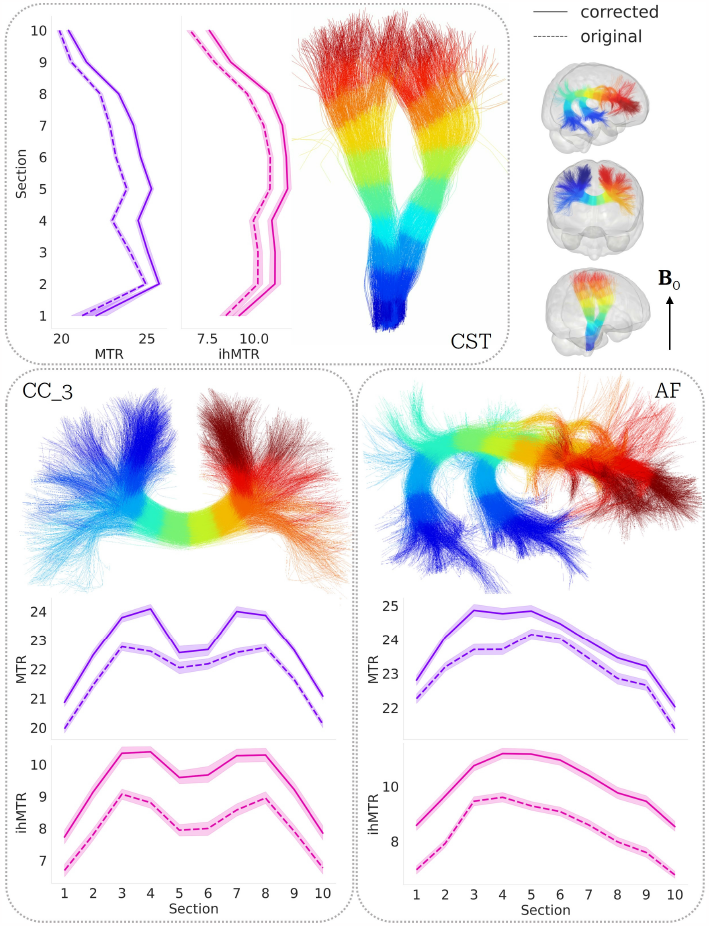
Track-profiles of the CST, CC 3 and AF, for original and corrected versions of MTR and ihMTR. Illustrations of the bundles, on which sections are color-coded, provide visual support for better understanding the profiles. These help locating the bundles in the brain, with the approximate direction of **B**_0_ also shown. Note that these illustrations are templates and do not correspond to the actual data. Moreover, the track-profiles are the means of every subjects and sessions and thus the colored shade around the profiles gives an idea of the variance.

## 4| DISCUSSION

### 4.1| Single-fiber characterization

The orientation dependence of MTR and ihMTR was successfully characterized using the method for singlefiber voxels, described in section 2.3. Indeed, the shapes of the orientation dependence curves presented on figure 3 and in Supporting Information correspond very well to the findings of Morris et al. ^19^. Moreover, the observed behavior of the voxel count could be explained in two parts. Firstly, it is mathematically expected that the number of fiber orientations separated by an angle *θ*_a_ of the main magnetic field is increasing with *θ*_a_. Indeed, the cone of possible orientations at a given angle becomes wider when the angle increases, in a manner similar to figure 1. Secondly, it is also possible that the anatomical disposition of fiber bundles in the brain favors orientations perpendicular to **B**_0_, rather than parallel to it. In Supporting Information, we try to understand further these curves by looking at the orientation dependence of the raw images used to compute the MTR and ihMTR.

#### 4.1.1| Localization of angle bins in the single-fiber selection mask

Upon visual inspection of figure 4, we notice that most of the single-fiber voxels are located in very few large WM bundles, namely the corticospinal tract and the corpus callosum. As the main magnetic field usually points somewhere around the z-axis of the images, it is expected that the principal projection fibers encompass most of the low angle bins for single-fiber voxels. This limitation for single-fiber voxels also explains why most of the high angle bins (40 to 90 degrees) are found in the central part of the CC, which is known to be a region where only one bundle passes through. The CC also contains voxels of lower angle bins, which are however shared with the CST.

#### 4.1.2| Corpus callosum versus whole white matter

Using the fortunate fact that the CC encompasses the whole range of angles from 0 to 90 degrees, we were able to compare the characterization of a singular structure with the one from the whole WM. Figure 5 confirms the hypothesis that the CST impacts the characterization of the whole WM at low angle bins. Indeed, the clear shift in MTR and ihMTR seen at low angles could indicate a high myelination of the CST. As shown previously on figure 4, the CST is composed mostly of voxels with angles spanning from 0 to 40 degrees. This concurs very well with the fact that the shift in measures means also occurs between these angles. It is also important to keep in mind that the CC shares most of its 0 to 30 degrees voxels with the end of the CST, meaning that the shift could be even more important in other parts of the brain. Moreover, both curves match above 40 degrees, which is to be expected. Indeed, as seen on figure 4, most of the high angle voxels are located in the CC, which means that both the whole WM and the CC characterization should be based on the same voxels at these angles. The presence of a few high angle regions that are not in the CC, along with the perfect match of the CC and whole WM curves for MTR, indicate similar macromolecule content between the CC and these other parts of the brain. However, the slight difference of the ihMTR trend from 80 to 90 degrees suggests that the myelin content might be lower in these regions of the CC than the regions highlighted in red at the bottom of the coronal slice on the first row of figure 4.

### 4.2| Crossing fiber characterization

Previous sections accessed the various subtleties of the orientation dependence of MT measures in single-fiber voxels. However, this ignores a large fraction of WM voxels, as crossing fibers are estimated to compose 60 to 90% of the total number of voxels in a typical brain ^22,23^. For this reason, we extended the typical single-fiber characterization methods ^16,19^ to be sensitive to multiple orientations in a single voxel, using fODFs instead of the average diffusion tensor from DTI.

The fact that the orientation dependence in voxels with two crossing fibers seems to follow a linear combination of two single-fiber characterization curves is encouraging from the perspective of understanding and correcting such dependence of MT measures. Indeed, the clear contribution of each orientation dependencies in the voxel suggests that correcting the measures for each fiber’s dependence would be feasible. This is also supported by the similar results obtained from voxels of three fibers crossing, which together with the single fibers and crossings of two fibers make up for the vast majority of WM. However, the behavior of the 3D curves when the fibers are orthogonal to each other and one is aligned with **B**_0_ is peculiar. We expect that the maximum measure would be seen when both fibers are oriented with an angle that gives the maximum measure for single-fiber voxels (around 90 degrees for MTR and 0 degrees for ihMTR). Although this case does produce a high measurement, the maximum is obtained in the previously described situation. We hypothesize that this is once again due to the higher myelination of the CST, as most of the voxels contributing to this situation come from the region where the CST is crossed by the MCP. These types of voxels are visible on figure 4 where one peak is in dark blue and the other is in dark red, and where both peaks share a similar fraction (blue-green color in the bottom part of the figure). This disqualifies the part of the centrum semiovale that would have otherwise counted, leaving the crossing of the CST and the MCP as the principal contributor to this particular case. This would explain also why ihMTR seems more affected by the phenomenon. Further research would be required to better understand this phenomenon and ultimately propose a correction method that takes this into account.

Another question raised from figure 6 is the shift in maximum amplitude observed between the measures for single-fiber and crossing fibers. For example, the maximum MTR for single-fiber characterization is slightly above 24.5, while the maximum for two fibers crossing, with fractions between 0.4 and 0.6, is around 23.0. Figure 7 allows for a better description of this phenomenon, as it shows that progressively adding a second fiber into a voxel slowly decreases this maximum measure. Furthermore, this figure also highlights a diminution of the maximum amplitude (*δ*m_max_) of the curves for MTR. This means that the separation between the maximum (at around 90 degrees) and the minimum (at around 40 degrees) values decreases, the curve flattens. While we do not have a theoretical explanation for this behavior, it is certainly something to consider for a correction method. Moreover, we observed on all the dataset a linear trend for the diminution of *δ*m_max_ with respect to the fraction of the dominant peak (**p**_1_). Fitting this trend then allows for a continuous description of the phenomenon, useful later for the correction method.

### 4.3| Correction results

The visual effects of the orientation dependence of MTR and ihMTR displayed on figure 8 are worrisome, especially since these measures are not known to have high contrasts. For instance, if one were to do tractometry on the corpus callosum, there would be a drop of MTR where the CC is oriented around 40 degrees with respect to **B**_0_. However, this variation of MTR is not due to a decrease of myelin content, rather the mere orientation of the WM fibers, which is problematic in the case of white matter integrity studies. Moreover, one could compare the track-profile of the CC and the CST, and see a major difference of myelination from ihMTR. Although this difference might be real, it is certainly amplified by the effect of the orientation dependence. Previously shown plots also demonstrate the striking variation of MTR and ihMTR with respect to the angle *θ*_a_. It is thus of great importance to be aware of this orientation dependence and try to correct the measures accordingly.

Therefore, we propose a correction method, presented in section 2.5, that is able to effectively remove the orientation dependence of MTR and ihMTR. As shown by figure 3, the corrected curves are almost free of variations along *θ*_a_, especially for MTR. The small variations of the corrected ihMTR curve correspond to angles where the polynomial fit does not perfectly match the data, due to the lower SNR of this measure. The method is also effective at reducing the orientation dependence in crossing fibers voxels, as illustrated by figures 6, 7 and even in the case of three fibers crossing. Although these results are not as convincing as for the single-fiber case, they still show curves that are much less dependent on orientation than the uncorrected ones. On figure 7, we see that corrected MTR measures tend to follow a flat trend, as the variation of MTR stays between a few decimals regardless of peak fraction. This suggest that our correction method rightfully accounts for the observed variation of maximum amplitude. As for ihMTR, which suffers from the scarcity of data from 0 to 20 degrees on figure 7, it is difficult to judge the correction results since the orientation dependence is relatively low at higher angles. Figure 6 provides more insight on ihMTR correction, as we observe the disappearance of the high ihMTR peak when both fibers are around 20 to 30 degrees, and the flattening of the majority of the surface. However, a large ihMTR spike remains when one fiber is parallel to **B**_0_ and both fibers are orthogonal. The same phenomenon is observed for MTR, which is nonetheless mostly composed of a flat surface after correction. This refers back to the previous observation of high MTR and ihMTR in such circumstances for the original measures, as discussed in section 4.2. Indeed, since this behavior is not present in the single-fiber characterization curves used as a reference for the correction, it causes over-correction. Nevertheless, both MTR and ihMTR corrections on crossing fibers appear successful elsewhere and the problematic situation still shows lower amplitude than the uncorrected measures. Furthermore, figure 8 demonstrates the ability of the correction method to visually remove the orientation dependence of MTR. Due to the noisier nature of ihMTR, the correction results are a little less appreciable.

The above discussion addressed the results of the correction method using the whole WM single-fiber characterization as a reference. On figure 5, we show that the use of a characterization limited to the corpus callosum does not properly correct the orientation dependence of the whole WM. Indeed, the observed decrease of the measures at low angles results in the over-correction of MTR and under-correction of ihMTR, which is also noticeable in the difference rows of figure 8. The CC method can correct the orientation dependence of the CC itself, but is not suitable for the whole WM on average. This, combined with the fact that the CC characterization differs from the whole WM characterization, suggest that a bundle specific approach could be useful. However, this becomes a much more complicated problem for crossing fibers, as bundles cross each other.

### 4.4| Impacts of correction on tractometry results

The work from Edde et al. ^28^ offers a unique opportunity to explore the effects of the proposed correction method on tractometry, namely bundle-average and track-profile results. In order for a correction method to be valid, it has to be able to keep the inherent trends of measurements while removing the false contribution of orientation. An inspection of the bundle-average results from figure 9 suggests that our method does not break the trends of MTR and ihMTR, as far as mean measurements in bundles are concerned. Indeed, the relationship between bundles seems to be roughly conserved, with the exception of the CST. This is the only bundle that follows more or less a single orientation throughout its length, meaning that the CST is the most susceptible bundle to be impacted by correction when compared to other bundles. Since the orientation of the CST with respect to **B**_0_ remains in the 0 to 20 degrees range, its MTR should overall increase slightly, while its ihMTR should barely change, as it is already at the maximum value. Figure 9 shows that the mean MTR of CST was originally similar to CC_6 and SLF_2, and lower than IFOF and ILF. After correction, it is now at the same level as IFOF and ILF, and higher than CC_6 and SLF_2. As for ihMTR, we observe a transition from CST being the highest bundle to CST being in the high-end part of the mean, which we believe to be more plausible. Indeed, both changes to MTR and ihMTR for the CST seem valid, as the CST would now be one of the highest myelinated bundle without being significantly over every bundles. Another important factor to consider is the fact that the correction process does not seem to induce variability, as demonstrated by the seemingly unchanged ranges of measures for each bundles.

Previous sections highlighted that MT measures can vary solely because of the orientation of the underlying WM fibers. This means that bundles of fibers should also be affected by this orientation dependence of MT. Track-profiles such as the ones presented on figure 10 provide a way of analysing various sections of a bundle, allowing the study of this effect. For instance, the almost unchanged trend of the CST profiles concurs with the constant orientation along the bundle. The overall increase of corrected MTR throughout the sections can be explained by the originally lower level of MTR at around 0 to 20 degrees, compared with the maximum level found from 80 to 90 degrees. However, the small increase in ihMTR, which is already supposed to be at a maximum level at these angles, hints at a slight tendency of the method to induce higher measurements. Bundles like the CC and the AF possess a wide variety of orientation angles and should thus be impacted by correction in various ways. Indeed, it is clear upon inspection of the CC_3 bundle and with knowledge of the previous characterizations that sections 5 and 6 (perpendicular to **B**_0_) are affected in a very different way by the orientation dependence compared to sections 1, 2, 9 and 10 (mostly parallel to **B**_0_), and more so to the remaining sections which are oriented somewhere between 20 and 70 degrees. This is effectively observed on figure 10 as the new corrected track-profiles are quite different from the original ones, especially for MTR. This shows a possibly more correct distribution of MTR along the CC 3, with the sections of crossing fibers (3, 4, 7, 8) having the highest values by far now. As for ihMTR, it is simply increased overall to match the maximum values from the characterization, except for the end-point sections which are indeed less increased. This is perhaps a bias of the whole-brain characterization method and the ihMTR should maybe not become that high. Similar effects can be observed on the AF bundle, as the regions perpendicular to **B**_0_ (5 to 10) present constant increase, while section 3 shows a good example of the opposite behavior of fibers with 0 to 20 degrees of orientation for MTR and ihMTR. Overall, it is difficult to attest that these changes are closer or not to the real distribution of MT measures. However, it is made clear that the original track-profiles are affected by the orientation dependence of MT and should be analysed carefully.

### 4.5| Limitations and future work

Despite providing encouraging results, the correction method still needs refinement in the way it handles crossing fibers. For the moment, the method does not properly deal with the high measures occurring when one fiber is parallel to **B**_0_ and the other is perpendicular to it, resulting in a over-correction. While this situation might not arise in a large number of voxels, it is still important to tackle. In the same vein, the correction of ihMTR in crossings is not as good as MTR, because we were not able to use the *f*_*δ*m_max (**p**_1_) factor taking into account the varying maximum amplitude. Another concern of the method is the difference in orientation dependence curves between WM structures, as highlighted throughout the manuscript and by Morris et al. ^19^. Indeed, the whole WM characterization approach used in this work might not be fully adequate for every single WM bundle around the brain. Our whole WM method might be fine on average throughout WM, but we are also probably falsely correcting certain regions. Moreover, tractometry showed how strongly the track-profiles can be impacted by the orientation dependence of MT, also suggesting that a bundle specific correction method could be interesting. Nonetheless, it is important to emphasize that not applying any correction is probably worst than using our correction method, even with its few flaws.

This variation of the orientation dependence between bundles brings the idea of bundle specific approaches for characterization and correction. Indeed, if each bundle had its own characterization curve, it would ensure that they are all accurately corrected. However, such method would require more development and reflection, as it would not be trivial to deal with crossing fibers. It might be possible to correct using a modeling algorithm such as COMMIT ^39^ retroactively. On another note, a priority should be to adjust the correction method to take into account the high measures appearing when one fiber is parallel to **B**_0_ and the other is perpendicular to it. This work also opens possibilities for future work unrelated to the limitations. For example, it would be interesting to explore the behavior and efficiency of the characterization and correction methods when used on MS patients. The characterization would probably have to be performed in a normal-appearing WM mask, free of lesions. Comparing the tractometry results for such patients with and without our correction method might uncover false assumptions about the myelination along certain tracts. Furthermore, the proposed methods are not limited to MT measures, as it can take any measures as input, such as T1-weighted images.

## 5| CONCLUSION

In this work, we first reproduced results from Morris et al. ^19^ regarding the orientation dependence of MT measures in single-fiber voxels. We also went a step further and explored the behavior of MT measures in crossing fibers voxels, showing that this orientation dependence follows roughly a linear combination of the single-fiber trend. To do so, we established characterization methods for both single and crossing fibers voxels. Using all this knowledge, we were able to come up with a correction method, effectively removing the orientation dependence of MT measures. We demonstrated the efficiency of our method both on the images directly and on the corrected plots. Furthermore, we showed the important impacts of the orientation dependence of MT measures on tractometry results. As a first attempt at understanding the shapes of the orientation dependence, we also showed the progression of this dependence throughout the calculation of the measures, starting from the raw images.

This work is a first attempt at providing a complete characterization of the orientation dependence of MT measures in all voxels of the WM, leading to the correction of said measures. As pointed out in the manuscript, the proposed method can still be refined, especially regarding the precise characterization of crossing fibers and the impact of different bundles. Nonetheless, our proposed method provides MTR, MTsat, ihMTR and ihMTsat measures that are globally free of orientation dependence.

## ACKNOWLEDGMENTS

The authors would like to thank everybody involved in the data collection from the past study ^28^, namely the researchers, the participants and the radiology team.

## Author contributions

Philipe Karan drafted the manuscript, designed and coded the characterization and correction methods, and contributed to analysing and interpreting the results. Manon Edde contributed to data collection (along with other collegues) and processing. Maxime Descoteaux and Guillaume Gilbert contributed to data collection and study design, and helped review the manuscript. Muhamed Barakovic and Stefano Magon contributed to the study design of the dataset. All authors revised the final version of the manuscript.

## SUPPORTING INFORMATION

Every plots presented in the results section of the manuscript are also available for all subjects and all sessions, following this link: https://doi.org/10.5281/zenodo.8383426. The actual data can be shared upon request to the authors. The Supporting Information is available as part of the online article.

