## Supporting Information for "Data-driven characterization and correction of the orientation dependence of magnetization transfer measures using diffusion MRI"

### S Supporting information

#### S.1 Variability of the characterization

The intra-subject variability can be assessed by comparing the characterization curves obtained from the multiple sessions within each individual subject. Then, a mean of theses sessions for each subject can allow for an inter-subject variability assessment, by comparing the means of the subjects. If these tests show very low variability of the characterization, a global correction approach might be conceivable. However, if the results show some degree of variability, as shown by Morris et al.[1], a subject-based approach would be better suited.

In addition to the orientation dependence of MTR and ihMTR presented by Morris et al.[1], we also characterize the saturation measures defined at equations 2 and 4, namely MTsat and ihMTsat. As these measures are supposed to be less impacted by $T_1$ relaxation, which is also known to have an orientation dependence[2], the characterization curves might be different from the ratio measures. Thus, a comparison of both ratio and saturation measures is necessary.

The methods presented in sections 2.3 and 2.5 depend on a few variable parameters, namely the FA threshold for the mask, the bin width and the tissue mask. Increasing the FA threshold would increase the anisotropy of the selected diffusion tensor, and reduce the number of selected voxels. The bin width impacts the smoothness of the produced plots, reducing noise whenever the amount of voxels is low. In this study, we evaluate the effect of various FA thresholds (0.5, 0.6, 0.7), bin width (1°, 3°, 5°, 10°) and the two previously discussed tissue masks, whole white matter and corpus callosum (CC).

Figure S.1 shows low variability of the orientation dependence intra and inter-subject. Indeed, while the curves have different shifts in MTR and ihMTR, they all follow a similar trend. We observe that most of the variations to the trend come from low angle bins (around 0 to 10 degrees), which have the lowest voxel count. This effect is also visible in figure 3. Although session 3 of subject 2 deviates from the other sessions, most subjects feature a degree of variability comparable to the ensemble of sessions 1, 2, 4 and 5 of subject 2. Thus, the first row of figure S.1 shows an example where a single session diverges from the mean.

Figure S.2 in the appendix demonstrates that the choice of bin width for single-fiber analysis solely impacts the visual aspect of the characterization and the number of voxel per bin. Indeed, curves created using bin widths of 1, 3, 5 or 10 degrees all look the same. However, this does not apply to crossing fibers characterization, as the low number of voxels per bin forces the use of wider bins (10 degrees). Moreover, figure S.3 in the appendix illustrates the effects of the FA threshold, as the curves have a slight gain in amplitude when the threshold is higher, and also seem to become

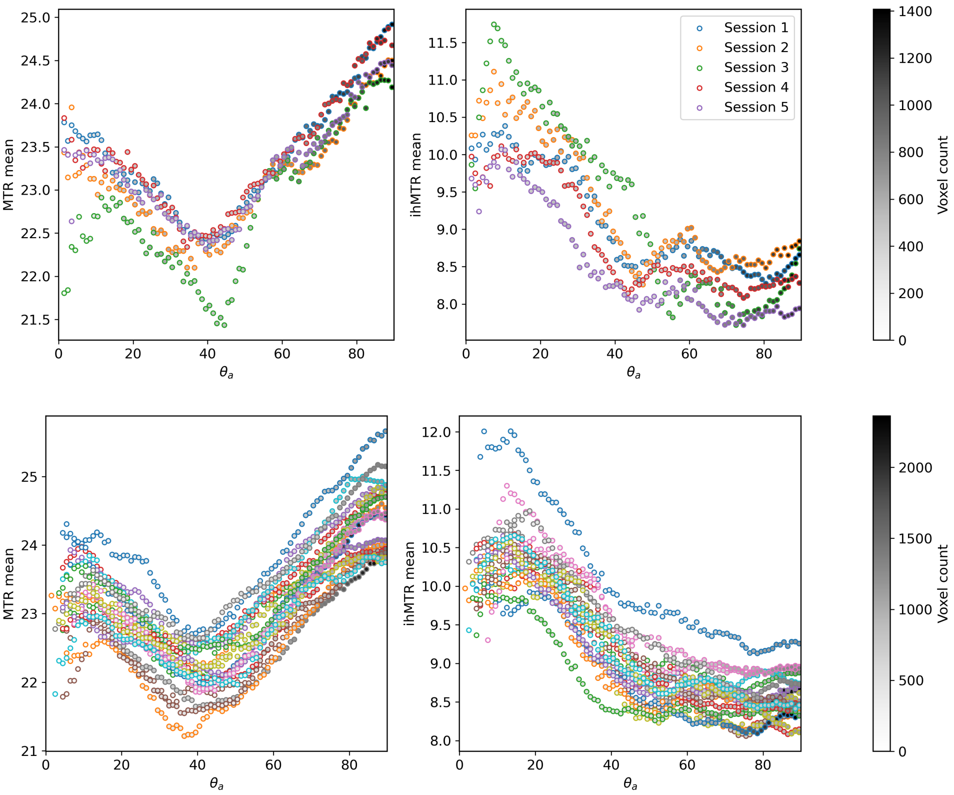


Figure S.1: First row: mean MTR and ihMTR with respect to the angle $\theta_a$ for the 5 sessions of subject 2. Second row: mean MTR and ihMTR with respect to the angle $\theta_a$ for the 20 subjects. For each subject, the 5 sessions were averaged to produce these results.

more noisy due to lower voxel count. We also show that saturation measures have an orientation dependence very similar to ratio measures, as presented in figure S.4 of the appendix.

Once again, the results of the characterization for the 20 subjects look very similar to the ones from Morris et al.[1]. The low variability observed on figure S.1 indicates that the orientation dependence of MTR and ihMTR is a precise phenomenon. The offset between the curves can be explained by the variations of myelin content (and non-myelin content in the case of MTR) across subjects. The positioning of the head inside the scanner can also be a major contributor to the variability of the orientation dependence, both across subjects and across sessions. Indeed, a slightly rotated head compared to a reference position will have small variations of the angle $\theta_a$ in voxels affected by the rotation, which ultimately affects the angle bins in which these voxels

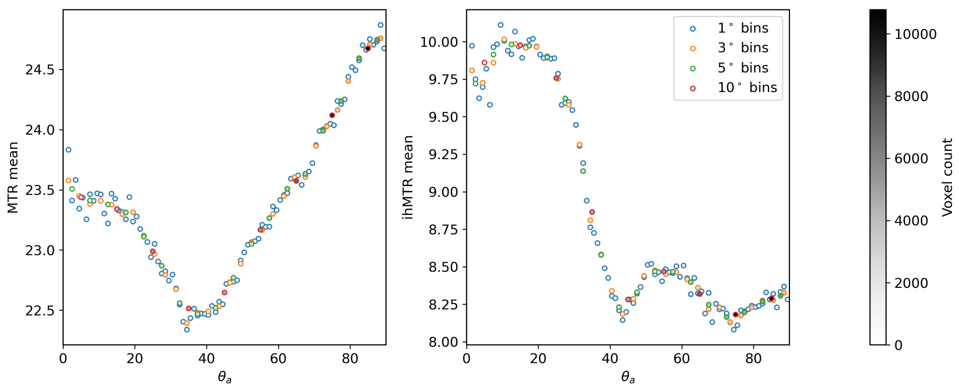


Figure S.2: Mean MTR and ihMTR with respect to the angle $\theta_\mathrm{a}$, for various bin widths.

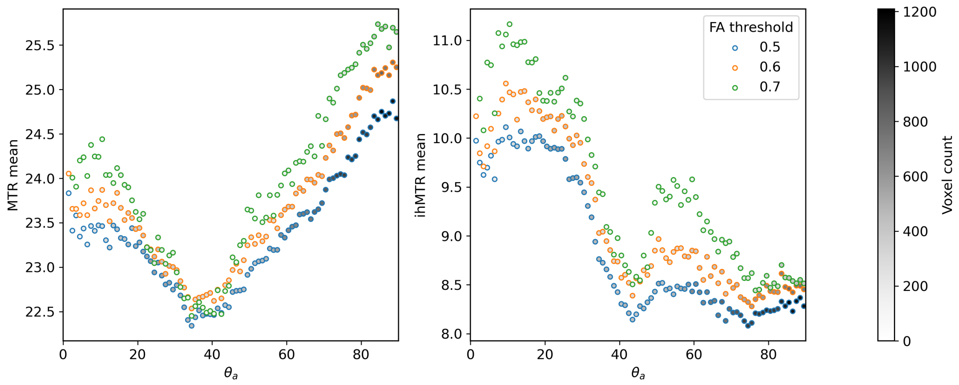


Figure S.3: Mean MTR and ihMTR with respect to the angle $\theta_\mathrm{a}$, for various FA thresholds.

are classified. Thus, some structures with an inherently higher myelin content might end up at different position on the curve, changing its shape. A good example of that would be if the orientation of the CST with respect to $\mathbf{B}_0$ changed between subjects or sessions. In that case, voxels that were contributing as high MTR and ihMTR for some angle bin would be counted in a different bin, effectively moving the low angle local maxima observed on the curves. The impacts of different WM bundles on the characterization method is discussed further in section 4.1.2. The lower voxel count at low angles can also explain some of the variation observed, as is shown by the stronger deviation from the polynomial fit at low angles on figure 3.

The fact that the curves from figure S.2 follow the same trend no matter the bin width indicates that bins of 1 degree width contain enough voxels to produce representative means, for single-fiber characterization. However, small variations are still visible at low angles and they decrease when the bins become wider. Regarding the effect of the FA threshold, the increase in MTR and ihMTR amplitude with higher FA can be attributed to an increase of myelin content. Indeed, a higher FA means that WM is more anisotropic, which is possibly correlated to an increase of WM myelination.

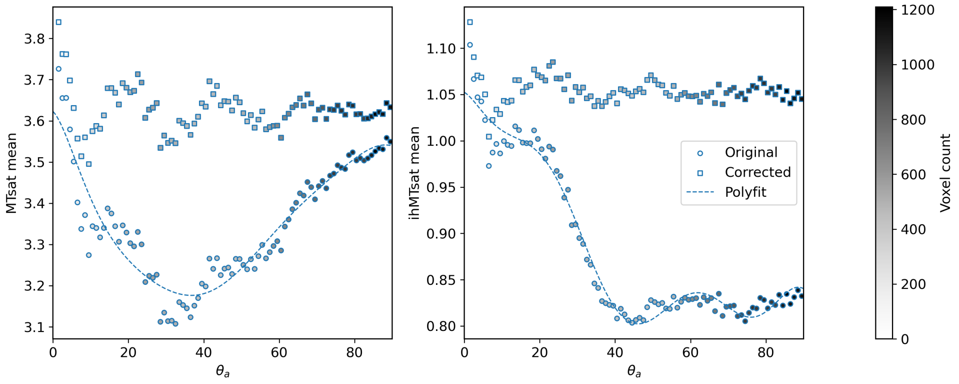


Figure S.4: Mean MTsat and ihMTsat with respect to the angle $\theta_a$.

Although figure S.4 illustrates that saturation measures (MTsat and ihMTsat) possess an orientation dependence that is overall similar to MTR and ihMTR, there are still notable differences, seen across the dataset. These differences most likely come from the $T_1$ correction of the saturation measures, as $T_1$ also has a known orientation dependence[3].

#### S.2 Understanding the orientation dependence shape

To better understand the shape of the orientation dependence curves produced, the single-fiber characterization method can also be applied to the raw images used to compute MTR and ihMTR. Indeed, studying the angular dependence of $S_+$, $S_-$, $S_{+-}$, $S_{-+}$ and $S_0$ could help clarify the shapes previously obtained and the contradictory results found[4,1].

Figure S.5 presents the evolution of the orientation dependence, from the raw images to the final measures. The first row shows the subtle differences between the orientation dependence of the mean single frequency-offset pre-pulse ($(S_+ + S_-)/2$) and the mean dual alternating frequency-offset pre-pulse ($(S_{+-} + S_{-+})/2$). It also highlights the presence of a small shift between the $S_+$ and the $S_-$ signals, which is due to an asymmetric effect of MT[5]. Meanwhile $S_{+-}$ and $S_{-+}$ share identical trends. The subtraction of the orange and green curve of the top-right plot gives the ihMTR mean, showed also in figure 3. The MTR mean is simply the inverse of this orange curve. The top-left plot also shows that dual alternating frequency-offset pre-pulses $S_{+-}$ and $S_{-+}$ have the exact same orientation dependence, while the single frequency-offset pre-pulses $S_{+}$ and $S_{-}$ present a small shift.

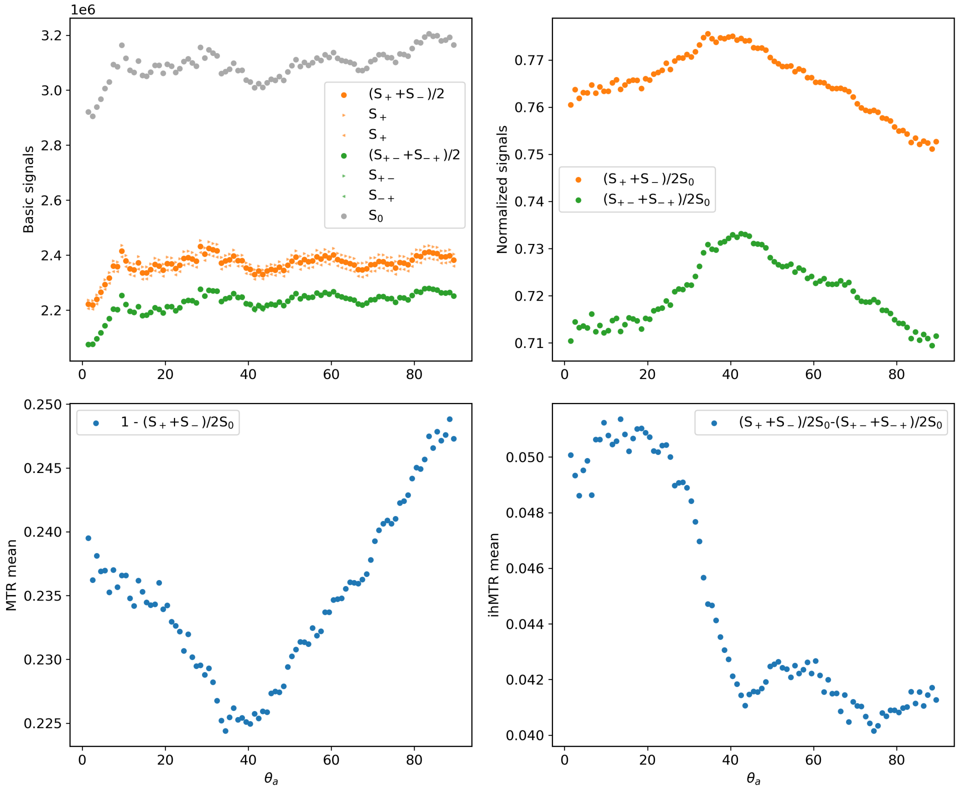


Figure S.5: First row: orientation dependence of the raw images (reference signal $S_0$, positive/negative/mean single frequency-offset pre-pulse and positive-negative/negative-positive/mean dual alternating frequency-offset pre-pulse) and their normalized versions. Second row: corresponding orientation dependence of MTR and ihMTR mean, equivalent to the original measures of figure 3.

With the present results of orientation dependence in single-fiber voxels supporting the findings of Morris et al.[1], the misalignment with the synthetic and simulated results from Morris et al.[4] is further confirmed. Indeed, the ihMTR orientation dependence shape simulated by Morris et al.[4] looks like the MTR behavior we observed, while it is completely different for the measured ihMTR. Figure S.5 demonstrates that both the normalized mean single frequency-offset pre-pulse ($(S_{+}$+$S_{-})/2S_0$) and the normalized mean dual alternating frequency-offset pre-pulse ($(S_{+-}$+$S_{-+})/2S_0$) have trends that are similar to the simulated one for ihMTR. The small differences between these two shapes lead to the measured ihMTR orientation dependence. Since the simulated dependence stems from a dipolar-like orientation dependence, the dual alternating frequency-offset pre-pulse, supposed to decouple the dipolar order, should not present this kind of trend. Results from figure S.5 suggest that some dipolar order still remains with the use of a dual pre-pulse, creating an ihMTR orientation dependence that does not follow the simulated one. Although the aim of this work is not to dwell into the physics of the orientation dependence shape, this glance at the behavior of raw images could be a first step towards understanding it.

#### S.3 Supplementary materials

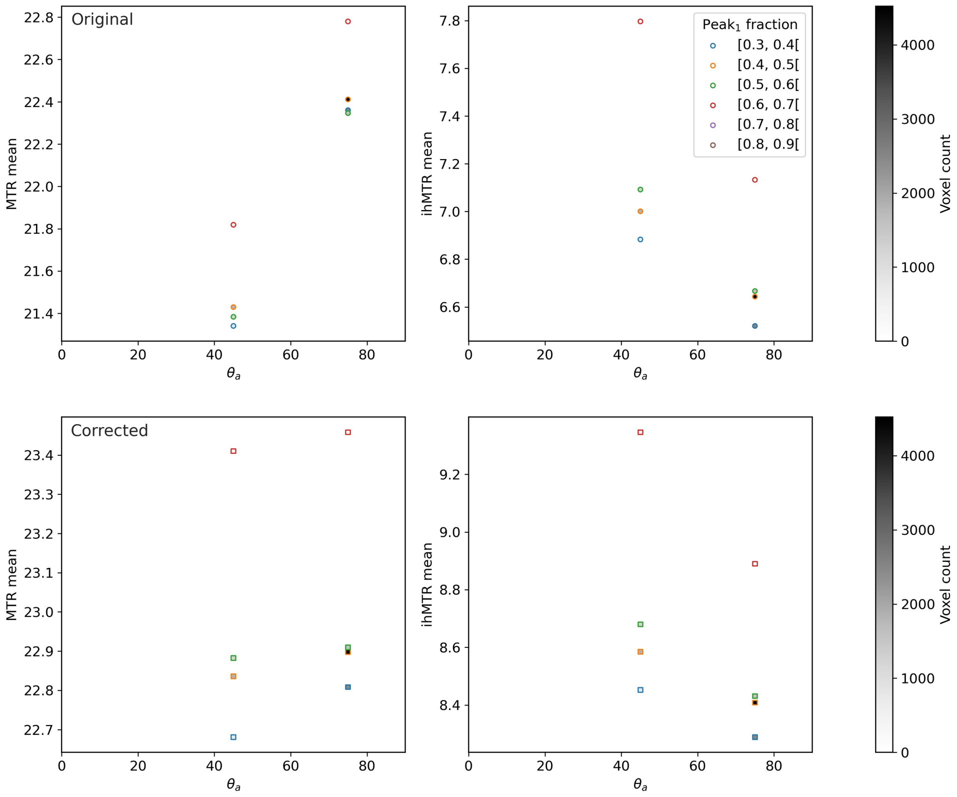


Figure S.6: First row: original mean MTR and ihMTR with respect to the angle $\theta_{\mathrm{a}}$, when angles $\theta_{\mathrm{a1}}$, $\theta_{\mathrm{a2}}$ and $\theta_{\mathrm{a3}}$ are equal, for various ranges peak fractions in the case of three crossing fibers. Second row: corrected mean MTR and ihMTR for the same situation as the first row.

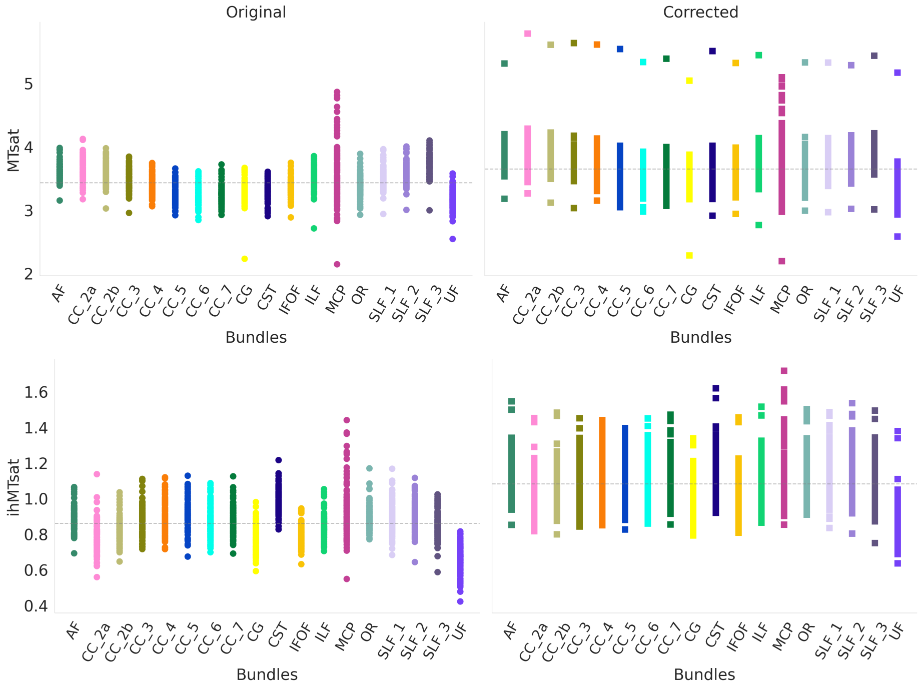


Figure S.7: Mean MTsat (top row) and ihMTsat (bottom row) of each selected bundles, for each subjects and each sessions. Again, the circle markers correspond to original measures (on the left), while the corrected measures are represented by square markers (on the right). The dashed grey lines represent the mean value of all the bundles, allowing for an easier comparison of changes between bundles. The left and right parts of bundles are averaged together.
